# Quantitative Analysis of Cytoplasmic Viscosity in Colorectal Cancer Cells by Differential Dynamic Microscopy of Genetically Encoded Nanoparticles

**DOI:** 10.1101/2025.11.13.687967

**Authors:** Steven Huysecom, Kiki Schouwstra, Samet Aytekin, Natalie Neyrinck, Yana Heyvaert, Boris Louis, Susana Rocha, Guillermo Solís-Fernandez

## Abstract

The viscosity of the cytoplasm plays a key role in regulating molecular diffusion and cellular mechanics, yet quantifying it in living cells remains technically challenging. Genetically encoded multimeric nanoparticles (GEMs) have emerged as powerful probes for intracellular microrheology; however, current analyses rely on single-particle tracking, which is limited by probe density, imaging noise, and expression variability.

Here, we combine GEMs with differential dynamic microscopy (DDM) to enable quantitative, non-invasive, and rapid measurement of intracellular viscosity using standard wide-field fluorescence imaging. DDM extracts particle dynamics from ensemble spatiotemporal intensity fluctuations, yielding reliable diffusion coefficients and viscosity values even in crowded or heterogeneous environments where tracking fails. Validation with fluorescent nanoparticles diffusing in water confirmed that DDM accurately reproduced theoretical viscosities across a wide range of particle sizes and concentrations. Comparison with single-particle tracking (SPT) demonstrated equivalent precision under dilute conditions and superior robustness under crowding.

To showcase the potential of this approach, we applied GEM-DDM to colorectal cancer cell lines with different metastatic potentials. Cytoplasmic viscosity correlated with aggressiveness, increasing from 1.9-2.3 cP in poorly metastatic to 3.6-3.7 cP in highly metastatic lines, consistent with greater macromolecular crowding and cytoplasmic reorganization reported in aggressive cells. Together, these results establish GEM-DDM as a fast, reproducible, and accessible platform for intracellular microrheology, providing new opportunities to link the physical state of the cytoplasm to cell function and disease progression.

**Statement of significance:** Physical properties such as cytoplasmic viscosity influence how molecules move and interact within cells, affecting metabolism, signalling, and disease progression. Measuring viscosity in living cells has been technically challenging and often invasive. Here, we introduce GEM-DDM as a quantitative, non-invasive image-based analysis method combining genetically encoded multimeric nanoparticles (GEMs) with differential dynamic microscopy (DDM) to measure intracellular viscosity using standard wide-field microscopy. We validate its accuracy against established single particle tracking (SPT) methods and demonstrate its biological relevance by showing that cytoplasmic viscosity increases with metastatic potential in cancer cells. This approach provides an accessible platform for studying how the physical state of cells influences their function and pathology.

## Introduction

Viscosity is a key regulator of biological processes, from molecular reactions to whole-cell behaviour. Within eukaryotic cells, the constant rearrangement of molecules between compartments depends on both passive diffusion and active transport, both of which are affected by cytoplasmic viscosity (Raja Venkatesh et al., 2023; Wilson et al., 2019). Changes in intracellular viscosity can therefore influence diffusion rates, reaction kinetics, and overall cellular dynamics. In red blood cells (RBCs), internal viscosity is essential for maintaining deformability and proper circulation (Ahmed et al., 2018; Mohandas et al., 1980; Renoux et al., 2019), with elevated cytosolic viscosity driving the membrane alterations observed in sickle-cell disease (Byun et al., 2012). Viscoelastic properties are not solely relevant for the behaviour of RBCs; they can also influence cell migration through dense extracellular environments. In cancer, where migration and invasion underpin metastatic progression, intracellular viscosity has been shown to inversely correlate with migratory capacity (Kole et al., 2005; Pan et al., 2022; Xiao et al., 2025).

Given its importance, a variety of strategies have been developed to probe intracellular viscosity, including fluorescent dyes such as sulforhodamine B (Srivastava & Krishnamoorthy, 1997) and viscosity-sensitive molecular rotors (Kuimova et al., 2009), passive nanoparticle tracers like gold particles (Guigas et al., 2007), and active microrheology techniques employing magnetically actuated nanowires or spin labels (Berret, 2015; Morse, 1986). Although these techniques have provided valuable insight, they typically require invasive delivery procedures that can perturb the cellular environment.

To overcome this limitation, methods based on the expression of genetically encoded fluorescent reporters were developed, allowing intracellular measurements without the need for probe delivery. For instance, fluctuation-based imaging techniques such as RICS and STICS extract diffusion coefficients from fluorescence intensity correlations in confocal image sequences, which can be related to local viscosity (Wiseman, 2013). While these approaches avoid physical manipulation of cells, they still require prior knowledge of the probe’s molecular radius and depend on confocal scanning microscopy, which limits their throughput and accessibility. More recently, Brillouin microscopy has emerged as a label-free tool for biomechanical characterization of cells (Keshmiri et al., 2023; Prevedel et al., 2019). However, widespread implementation of Brillouin microscopy has been hampered by the absence of commercially available systems and the need for a Brillouin spectrometer and confocal pinhole alignment. Together, these limitations highlight the need for non-invasive, reproducible, and easily accessible techniques to quantify intracellular viscosity in living cells. Genetically Encoded Multimeric nanoparticles (GEMs) have emerged as a breakthrough tool for intracellular microrheology (Delarue et al., 2018; Hernandez et al., 2024). These probes are built from self-assembling scaffold proteins fused to a fluorescent protein, that form bright nanoparticles of precise and uniform dimensions of approximately 20 or 40 nm in diameter. They can be expressed directly inside cells by transient or stable transfection, avoiding complications from exogenous nanoparticles such as inefficient cytoplasmic delivery, vesicle entrapment, or aggregation. Moreover, because this direct expression in the cytoplasm, they assemble spontaneously in transfected cells, avoiding invasive delivery steps. Importantly, GEM expression allows long-term monitoring of viscosity changes over time with minimal cellular stress. Their defined nanoscale architecture and size make them reliable physical tracers for characterizing intracellular dynamics (Hernandez et al., 2024). Despite these advantages, quantitative analysis of GEM dynamics has relied on single-particle tracking (SPT), which becomes error-prone in crowded or highly viscous environments where trajectories overlap (Korunova et al., 2024). To overcome this limitation, we implemented Differential Dynamic Microscopy (DDM) as a complementary, ensemble-based image analysis strategy to retrieve viscosity values from GEM dynamics. DDM extracts dynamic information from temporal intensity fluctuations in standard wide-field image sequences, allowing robust quantification of particle motion in both crowded and dilute environments (Cerbino & Trappe, 2008). By integrating GEMs with DDM into a unified framework, hereafter referred to as GEM-DDM, we established a non-invasive, accessible, and statistically powerful approach to measure intracellular viscosity directly from wide-field fluorescence data.

We validate GEM-DDM by benchmarking DDM-derived viscosity values against SPT measurements *in vitro* and apply it to a panel of colorectal cancer cell lines, revealing clear differences in cytoplasmic viscosity that correlate with metastatic potential. Beyond its technical advantages, this approach opens new possibilities for long-term, minimally invasive, and high-throughput studies of intracellular mechanics in living systems, offering a practical route to explore how biophysical properties such as viscosity evolve across cell states, differentiation, and disease progression.

## Results & Discussion

### Differential Dynamic Microscopy for Quantitative Microrheology

To quantify particle motion and extract microrheological parameters from wide-field fluorescence image sequences, we used DDM, an image-based analysis technique that extracts dynamics from spatiotemporal intensity fluctuations (Cerbino & Cicuta, 2017; Cerbino & Trappe, 2008; Verwei et al., 2022) (**Fig. 1A**). For each time lag τ, differential images were calculated by subtracting the image at time *t* from the image at time *t* + *τ*, removing static features and suppressing background noise. The Fourier transform of each differential image was calculated, and all transforms corresponding to the same time lag (*τ*) were averaged to obtain a mean Fourier image for that time lag. By radially averaging the mean Fourier image, we separated the dynamics at different spatial frequencies (wave vectors, *q*), obtaining an intensity amplitude for each *q*. The intermediate scattering function (ISF) was then calculated by plotting these amplitudes as a function of the corresponding time lag (*τ*). The ISF describes how spatial correlations decay over time: faster or steeper saturation indicates more rapid particle dynamics at the length scale of the corresponding *q*-vector.

**Figure 1:**
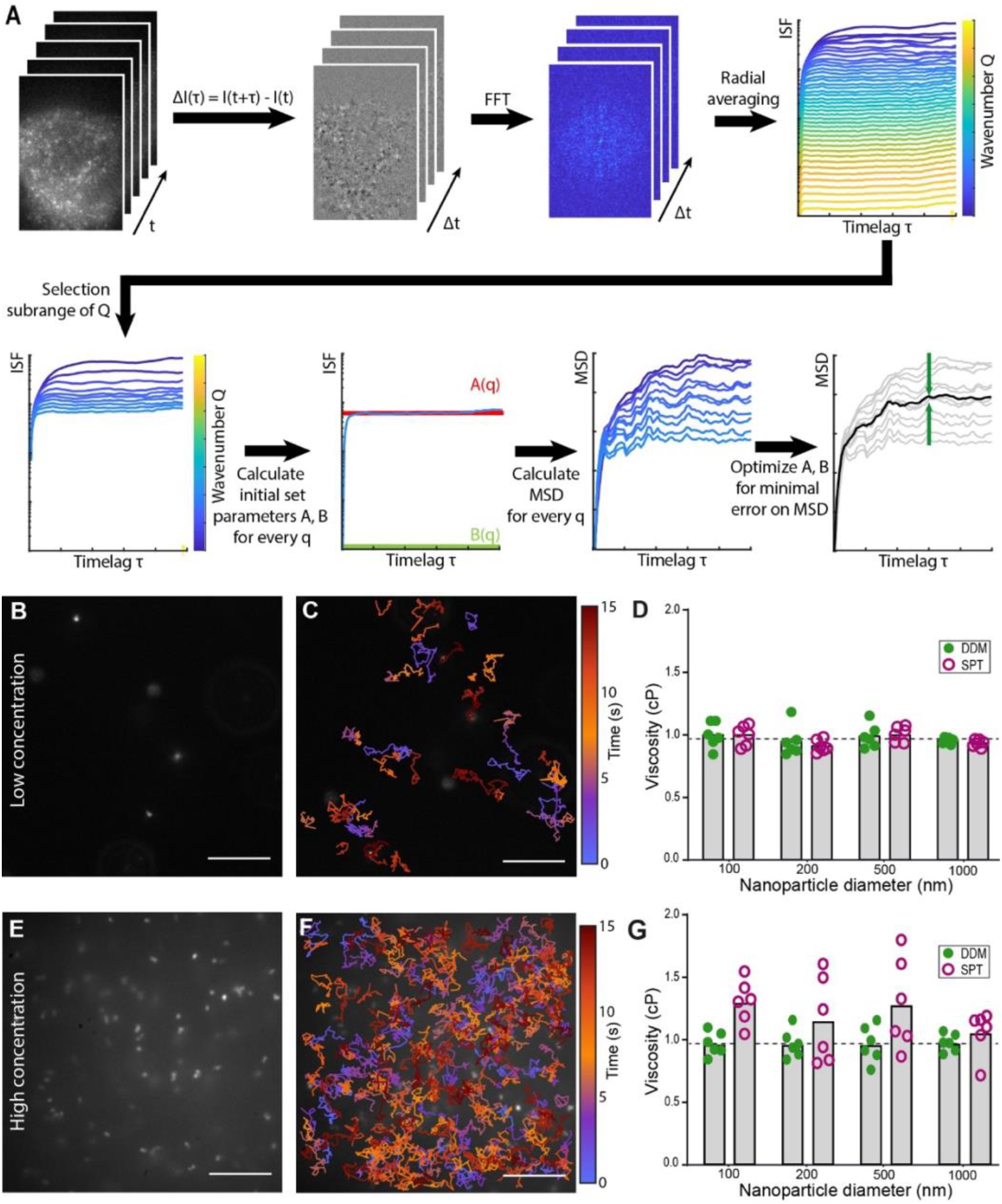
Overview of the analysis workflow of DDM and its validation by comparison with single particle tracking (SPT) of nanoparticles diffusing in water. : (A) The DDM-based analysis involves two steps with (i) the conventional DDM algorithm: calculating differential images and their Fourier transform for every timelag, and radial averaging to obtain the image structure function (ISF) per wavevector q. In an extension, a subset of q is selected to calculate the average mean-squared displacement of all particles, by an optimisation loop taking the saturation level A and baseline B of every ISF as optimising parameters. Partially adapted from (Edera et al., 2017). (B,E) Representative snapshot of 100 nm particles diffusing in water at diluted and crowed conditions. (C,F) Traces are labelled by timepoint – only traces suitable for microrheology analysis (containing more than 20 datapoints) were shown. (D,G) Measured viscosities for all samples in both diluted and crowded conditions, compared between DDM and SPT. Dashed black line gives the reference value of 0.932 cP for the viscosity of water at 23°C. All scalebars 10 µm.

For the analysis of both water and cytoplasmic viscosity, we selected *q*-vectors in the range 3–10 µm^−1^. These values correspond to spatial scales from about 600 nm, which reflects the diffraction limit and the displacement of the fluorescent nanoparticles within the exposure time, up to roughly 6 µm, comparable to the size of small cells.

Because intracellular diffusion dynamics can be complex (anomalous, Brownian, active, or confined), one cannot predict a single diffusion model that would adequately fit all ISFs. Instead, the algorithm was extended to convert each selected ISF into a mean-squared displacement (MSD) curve (Edera et al., 2017). First, the saturation level (A) and background (B) were estimated for each ISF. Then, using these initial set of parameters, each ISF was transformed into an MSD. An iterative optimization loop refined A and B to minimize the error across the set, producing a converged, robust average MSD. From the converged MSD, effective diffusion coefficients were obtained by linear fitting of the initial time lags, and viscosity was then calculated using the generalized Stokes-Einstein relation, assuming a predominantly viscous environment. In complex cellular environments where multiple diffusion regimes coexist, such as in the case of GEMs exhibiting both Brownian and directed motion, the overall MSD reflects this mixture through an anomalous profile: it may grow superlinearly before reaching a plateau, capturing not only effective diffusion and viscosity but also signatures of confinement and viscoelasticity. Although this study focused on the viscous regime, MSD analysis can, in principle, be extended to extract elastic and viscoelastic properties of the medium for more detailed microrheological characterization (Edera et al., 2017).

To validate the performance of our DDM algorithm and ensure accurate retrieval of diffusion coefficients and viscosities, benchmark experiments on fluorescent polystyrene nanoparticles diffusing in water were performed. Time-lapse images of fluorescent polystyrene nanoparticles with diameters of 100, 200, 500, and 1000 nm were recorded at low (< 0.0005 mg of particles per mL) and high concentrations (0.01 mg of particles/mL) to evaluate the effect of crowding on DDM performance (**Fig. 1B,E**).

Across all particle sizes and concentrations, DDM provided viscosity values in close agreement with the theoretical viscosity of water (0.932 cP at 23 °C). At low concentrations, the measured viscosities for 100, 200, 500, and 1000 nm nanoparticles were 0.98 ± 0.09, 0.93 ± 0.10, 0.97 ± 0.08, and 1.00 ± 0.19 cP, respectively. At higher concentrations, the values obtained were similar (0.97 ± 0.08, 0.97 ± 0.09, 0.96 ± 0.12, and 0.97 ± 0.06 cP for 100, 200, 500, and 1000 nm nanoparticles, respectively). These results demonstrate that DDM yields consistent results independent of particle size or sample crowding (**Fig. 1D,G**). This robustness arises from the ensemble-based nature of DDM, which analyses spatiotemporal intensity fluctuations rather than individual trajectories and is therefore unaffected by particle overlap or background heterogeneity. To benchmark these results, we compared DDM with conventional single-particle tracking (SPT). At low particle concentrations, SPT produced viscosities of 0.98 ± 0.07, 0.89 ± 0.04, 1.12 ± 0.04, and 0.84 ± 0.07 cP for 100, 200, 500, and 1000 nm nanoparticles, respectively – values that closely match both the DDM measurements and the theoretical viscosity of water (**Fig. 1D**). However, at higher concentrations, where particle density and background heterogeneity increase, SPT performance degraded substantially, yielding less reliable viscosities of 1.30 ± 0.19, 1.15 ± 0.29, 1.28 ± 0.31, and 0.06 ± 0.15 cP for the same particle sizes (**Fig. 1G**). Trajectories became shorter and less reliable due to overlapping paths and loss of particle detection, particularly for smaller particles that exhibited lower brightness and higher positional uncertainty. As a result, SPT overestimated viscosity values, while DDM continued to provide stable and accurate measurements under the same conditions.

Together, these validation experiments demonstrate that DDM provides accurate and reproducible viscosity measurements across a wide range of particle concentrations (0.01 to 0.0005 w/v%). Under dilute conditions, DDM achieves the same precision as SPT, while at high densities it remains reliable when SPT fails. This robustness stems from the ensemble-averaging nature of DDM, which ensures reproducibility even in heterogeneous or crowded systems where single-particle detection becomes unreliable. These results establish DDM as a robust, calibration-free method for quantitative microrheology, bridging the gap between single-particle tracking and more complex correlation-based approaches such as RICS or fluorescence correlation spectroscopy.

### GEM-DDM microrheology for cytoplasmic viscosity

Given the robustness of DDM, we applied this method to living cells to measure cytoplasmic viscosity using GEMs. To minimize interference from cellular autofluorescence, we developed a red-shifted GEM variant in which the original YFP was replaced with the red fluorescent protein mScarlet-I. This fluorophore was selected for its favourable spectroscopic properties and rapid maturation, providing bright and stable emission suitable for wide-field imaging. The resulting 40 nm GEM construct self-assembles into fluorescent nanoparticles following transfection and expression in cells (**Fig. 2A**). Cells expressing the red GEMs were imaged on a home-built wide-field microscope with a 561 nm excitation and an EM-CCD camera at 30Hz.

**Figure 2:**
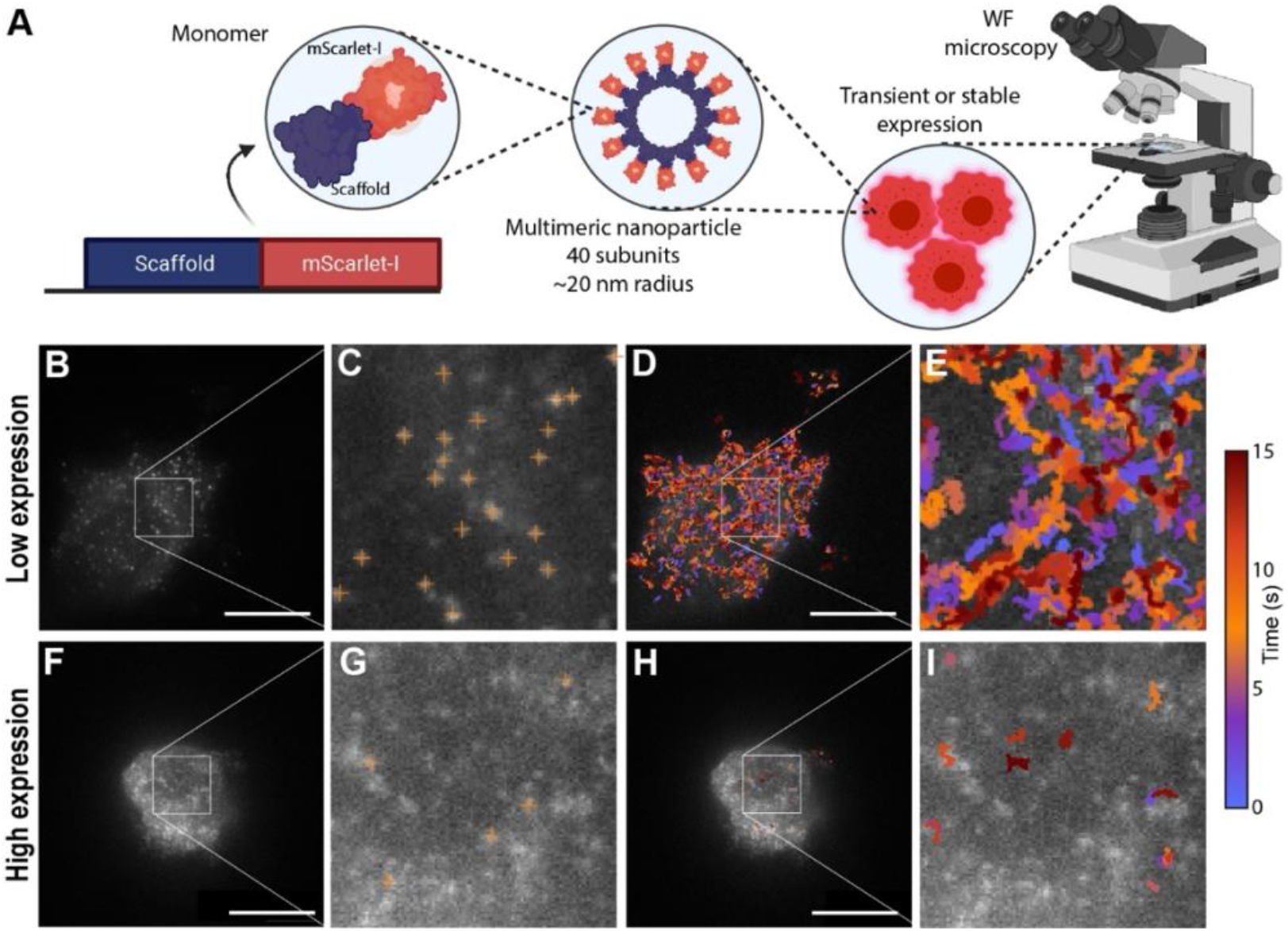
Evaluation of GEM-DDM workflow to measure cell viscosity benchmarked to SPT: (A) Experimental workflow involves the expression of GEM-mScarlet gene inside the cell, after which the nanoparticles form by self-assembly, and the fluorescent cell can be imaged with conventional wide-field microscopy. Partially adapted from (Shu et al., 2021). (B) KM12SM cell with low expression level with (C) zoom showing the localised particles in frame 1. (D/E) Low expression levels lead to many spread-out trajectories. (F) Highly expressing KM12SM cell with (G) zoom showing the localised particles in frame 1, showing many particles are missed due to heterogeneo background. (H/I) High expression levels biased the detection to a few very confined trajectories. All scalebars 10 µm

The performance of GEM-DDM in the intracellular environment was first evaluated by comparison with single-particle tracking SPT in individual KM12SM cells expressing GEMs at different levels (**Fig. 2B,F**). In cells with low GEM expression (**Fig. 3B,C**), individual particles were well resolved, which resulted in numerous and long trajectories distributed throughout the cytoplasm (**Fig. 3D,E**). However, only 3 out of 24 cells yielded SPT-derived viscosities within 10 % of the corresponding DDM values, indicating that SPT rapidly lost accuracy as particle density increased. Nonetheless, for this small subset, both methods yielded comparable viscosity values, with 2.51 cP on average measured by SPT and 2.57 cP by DDM. In the other 21 cells with higher GEM expression (**Fig. 3F,G**), only the brightest and most confined particles could be detected (**Fig. 3H,I**), leading to an overestimation of viscosity by SPT (5.86 cP) compared with DDM (2.22 cP). In conclusion, SPT showed to be less reliable in crowded cytoplasmic environments.

**Figure 3:**
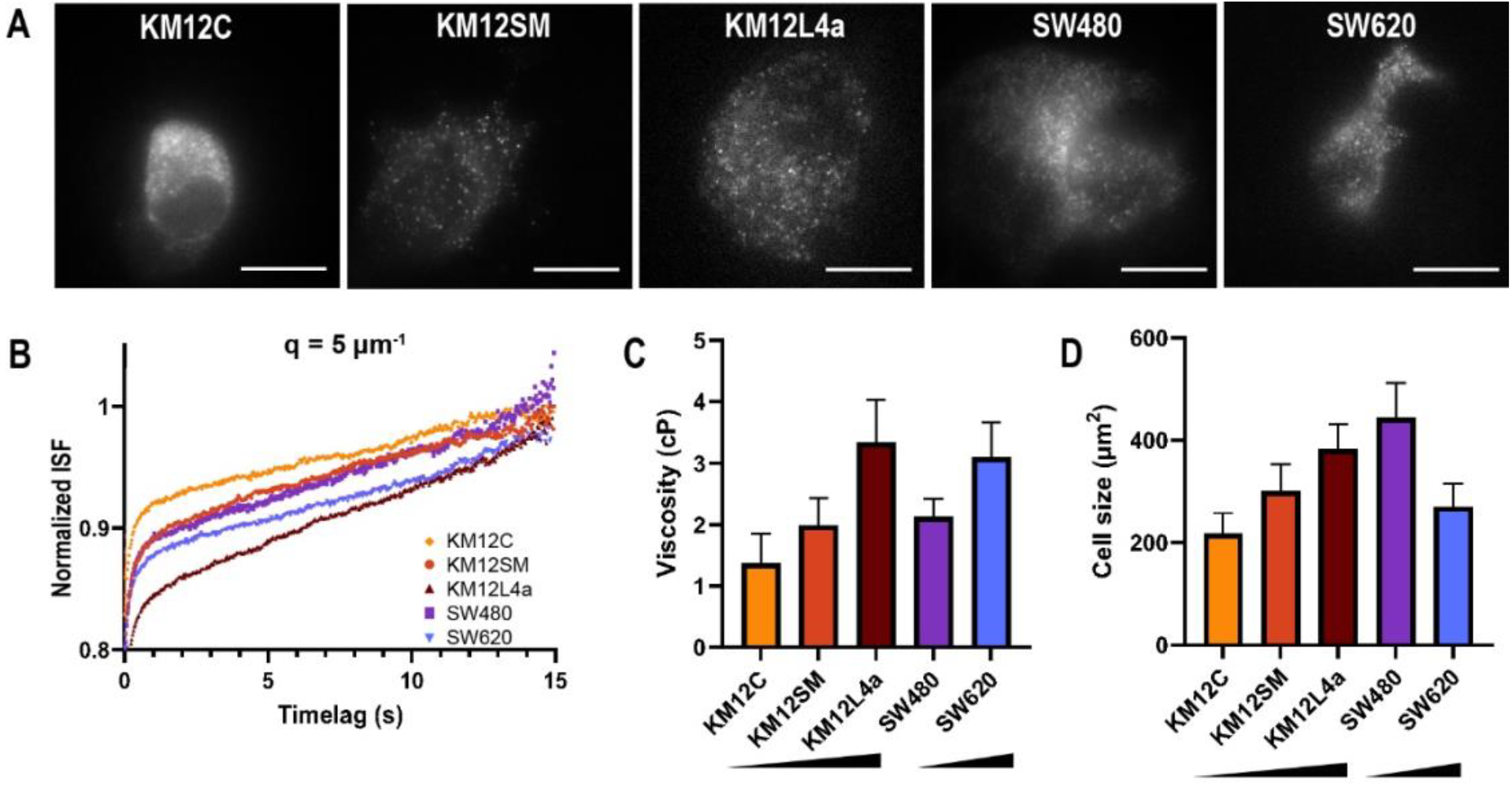
Measuring in vivo cytoplasmic cell viscosity with GEM probes and DDM based analysis. (A) Snapshots of representative cells per cell line. All show substantial crowdedness and differences in size. (B) ISF at wavevector q = 5 µm^−1^. The difference in dynamics can between non-/low-metastatic (KM12C, KM12SM, SW480) and highly metastatic cells (KM12L4a, SW620) becomes clear, showing a slower rise and thus dynamics for more metastatic cells. (C) Average viscosity per cell line. Plotted as geometric mean with 95% confidence interval, as all datasets showed a lognormal distribution. Number of cells n = 25, 24, 27, 28, 24 for KM12C, KM12SM, KM12L4a, SW480 and SW620 respectively. Black triangles indicate the increase of metastatic potential among cell lines. (D) Average size per cell line. Plotted as geometric mean with 95% confidence interval, as all datasets showed a lognormal distribution. Black triangles indicate the increase of metastatic potential among cell lines. All scalebars 10 µm

These results demonstrate that GEM-DDM provides a rapid, non-invasive, and broadly accessible approach for intracellular rheology. By analysing fluorescence fluctuations across multiple length scales and avoiding single-particle tracking, the method remains robust in crowding and heterogeneous environments, enabling reproducible viscosity measurements with standard wide-field fluorescence microscopy. The use of a single-color, genetically encoded probe eliminates the need for particle delivery or complex optical setups, simplifying the implementation of GEM-DDM and reducing perturbation to cell physiology. Together, these advantages establish GEM-DDM as a powerful and versatile platform that lowers the experimental barrier to intracellular microrheology and opens possibilities for systematic, high-throughput assessment of cytoplasmic viscosity in living cells.

### GEM-DDM analysis of Cytoplasmic Viscosity in Colorectal Cancer Cells

To demonstrate the applicability of GEM-DDM to physiologically relevant systems, cytoplasmic viscosity was quantified in a panel of colorectal cancer cell lines differing in metastatic potential. **Figure 3A** depicts representative fluorescence images showing dense GEMs distributions within the cytoplasm in all lines. This complex and crowded signal provided a suitable basis for quantitative analysis using DDM. Qualitative evaluation of the intermediate scattering functions (ISFs) extracted at a representative wavenumber of *q* = 5 µm^−1^ revealed distinct behaviours among the cell lines (**Fig. 3B**). In the poorly metastatic KM12C, KM12SM, and SW480 lines, the ISFs decayed rapidly, indicating fast decorrelation of particle positions and thus low cytoplasmic viscosity. Conversely, the highly metastatic KM12L4a and SW620 cells exhibited slower ISF relaxation, reflecting reduced particle mobility and higher viscosity.

Quantitative DDM analysis confirmed these differences (**Fig. 3C**). Within the KM12 family, viscosity increased with metastatic potential: 1.93 cP for KM12C (poorly metastatic), 2.22 cP for KM12SM (intermediate), and 3.72 cP for KM12L4a (highly metastatic). A similar trend was obtained for the SW family, with SW480 (poorly metastatic) showing 2.35 cP and SW620 (highly metastatic) reaching 3.58 cP. To assess whether these viscosity differences could be attributed to cell morphology, projected cell areas were measured (**Fig. 3D**). In the KM12 series, cell area increased with metastatic potential (239 µm^2^ for KM12C, 323 µm^2^ for KM12SM, 401 µm^2^ for KM12L4a), whereas the opposite trend was observed for the SW series (SW480: 463 µm^2^; SW620: 265 µm^2^). Even after normalization for cell size (**Supplementary Fig. S1**), highly metastatic cells consistently displayed higher cytoplasmic viscosity

The observed correlation between cytoplasmic viscosity and metastatic potential is consistent with prior reports linking increased macromolecular crowding, cytoskeletal remodelling, and altered metabolic activity to cancer aggressiveness (Aseervatham, 2020; Datta et al., 2021; Dessard et al., 2024).

Elevated viscosity may thus reflect a more crosslinked, densely packed cytoplasmic architecture in metastatic cells, potentially driven by enhanced cytoskeletal tension, accumulation of macromolecular complexes, or altered organelle distribution.

Together, these findings demonstrate the potential of GEM-DDM as a versatile and accessible tool for intracellular microrheology, enabling quantitative analysis of cytoplasmic viscosity across diverse cellular and physiological contexts. In addition to quantifying viscosity, this method provides a platform for future investigations into the interplay between intracellular mechanics, metabolism, and cancer progression.

## Conclusion

This study establishes GEM-DDM as a robust and accessible framework for quantitative intracellular microrheology. By integrating genetically encoded fluorescent nanoparticles with differential dynamic microscopy, this method overcomes the limitations of single-particle tracking and enables reproducible viscosity measurements in living cells using standard wide-field fluorescence imaging.

Beyond the methodological innovation, GEM-DDM provides a practical means to link cell mechanics with biological state. The observed correlation between cytoplasmic viscosity and metastatic potential underscores the value of viscosity as a biophysical marker of cellular organization and function.

The versatility and non-invasive nature of GEM-DDM make it well-suited for large-scale or longitudinal studies, offering a way to explore how intracellular physical properties evolve during disease progression, therapy response, or cellular differentiation. By enabling robust and accessible intracellular microrheology, GEM-DDM lays the groundwork for a broader mechanobiological understanding of cell physiology.

## Acknowledgments

We acknowledge additional financial support from Research Foundation of Flanders (FWO) research grant (G0C2422N), postdoctoral fellowships (GS: 12AML24N, for BL: 12AGZ24N), PhD fellowships (for SH: 11A0S25N, for SA: 1S95125N) and from the KU Leuven (IDN/20/021 and C14/22/085). YH and SR acknowledge the Global PhD partnership program between KU Leuven and Melbourne University (GPUM/24/010).

## Materials & Methods

### Nanoparticle Sample Preparation

Polystyrene nanoparticles (ThermoFisher FluoSpheres™ Size Kit #1, carboxylate-modified, red fluorescent 580/605 nm, 2% solids) were used for validation experiments. Particle suspensions were diluted in MilliQ water at either 1:1000 (low concentration) or 1:100 (high concentration) and vortexed for 5 min. A 10 µL drop was dropcasted on ozonated glass coverslips and sealed with a 120 µm thick, 9-mm diameter imaging spacer (Grace Bio-Labs SecureSeal™) and a second ozonated coverslip.

### Plasmid cloning

The original vector pCDNA3.1-pCMV-PfV-GS-Sapphire was a gift from Liam Holt (Addgene plasmid #116933; http://n2t.net/addgene:116933; RRID: Addgene 116933). Into this backbone we subcloned mScarlet-I and removed Sapphire through restriction enzyme cloning digesting with PacI and BsrGI.

### Cell Culture and Transfection

Colorectal cancer cell lines (KM12C, KM12L4a, KM12SM, SW480, SW620) were cultured at 37°C under 5% CO_2_ in DMEM supplemented with 10% FBS, 1% L-glutamax, and 0.1% gentamicin. KM12C and SW480 are poorly metastatic and derived from primary tumors, whereas KM12SM, KM12L4a, and SW620 are highly metastatic and derived from liver, lung, or lymph node metastases respectively (Camps et al., 2004; Hewitt et al., n.d.)(Camps, Hewitt). Cells were passaged at 80–90% confluency using trypsin-EDTA. For experiments, 5×10^5^ cells were seeded over the whole area of the 35 mm glass bottom dish (CellVis). Transfection of GEM plasmids was performed with FuGene HD (3 µL per 1000 ng DNA in 100 µL DMEM) directly on the gels after seeding the cells.

### Hydrogel Preparation and Glass Coating

Polyacrylamide hydrogels (37 kPa) were prepared following Cresens et al. with 4% w/v acrylamide, 0.17% w/v bisacrylamide, 0.25% w/v APS, and 0.25% v/v TEMED. 15 µL of these gels were cast in 35 mm glass-bottom dishes (20 mm micro-wells Cellvis), threated with bind-silane inside a 120 µm thick, 9-mm diameter wide imaging spacer (Grace Bio-Labs SecureSeal™). A Sigmacote-treated coverslip was added on top to maintain gel flatness during polymerization, which was performed at 37°C for 30 min, followed by overnight swelling at 37°C in MilliQ water. Hydrogels were functionalized with sulfo-SANPAH under UV light and coated with collagen for 2 h at 37°C.

### Fluorescence Imaging

Samples were imaged on a custom-built widefield microscope equipped with a 561 nm laser line 48 hours after transfection. A water-immersion objective (60x, Olympus, long working distance, 1.20 NA) was used to reach cells on top of the hydrogel. The pixel size was set at 107 nm per pixel. Fluorescence emission was filtered through a 532 nm long pass dichroic mirror and a 595/40 nm bandpass filter. Images were acquired on a EM-CCD camera in EM mode with adjustable gain, using 30.5 ms exposure and 500 frames per acquisition.

### Single Particle Tracking (SPT) Analysis

Nanoparticle trajectories were obtained and analysed using custom MATLAB code (Louis et al., 2020 – github Steven Huysecom). Particle candidates were identified through a Gaussian scanning approach and further localised using a phasor-based super resolution fit. Trajectories required a minimum of 50 steps, a maximum frame gap of 3, and an exclusion radius of 1500 nm. Mean-squared displacements (MSDs) were calculated for each trajectory, from which diffusion coefficients and viscosities were determined, as will be discussed in the next paragraph.

### Differential Dynamic Microscopy (DDM) Analysis

DDM analysis was implemented based on **Edera et al. (2017)** using custom MATLAB scripts and is depicted in **Fig.1B**. Preprocessing included rough segmentation of the cell from which further cell size was calculated and bleaching correction by correcting the average intensity of the frames over time. Differential images were generated by subtracting each possible pair of frames and each of them then Fourier-transformed. The Fourier images were averaged per timelag and radially averaged to construct image structure functions (ISFs) for each q-vector, ranging from the diffraction limit (≈ 233 nm) to the full image size.

For MSD calculations, a subset of wavevectors (1-10 µm^−1^) was selected to capture particle motion in the relevant range. The upper limit (q = 10 µm^−1^) corresponds to image frequencies of ∼623 nm, accounting for the diffraction-limited particle size and movement during the 30.5 ms exposure time. The lower limit (q = 1 µm^−1^) corresponds to image frequencies of ∼6 µm, which is big enough to capture the full movement inside a cell. Each wavevector has a corresponding ISF, from which baseline (*B(q)*) and saturation level (*A(q)*) are initially determined by

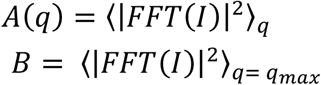

These values provide starting lists of *B(q)* and *A(q)* for the optimization procedure.

From each ISF, the mean-squared displacement (MSD) is calculated using the relation (**Edera et al. 2017**):

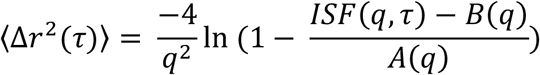

This results in a set of MSDs with some variance across q-vectors. An optimisation loop, implemented with MATLAB’s fminsearch, adjusts the *A(q)* and *B(q)* parameters to minimise the standard deviation of the MSDs. The optimisation uses a maximum of 10,000 iterations and a convergence cutoff of 10^−4^ on the MSD standard deviation.

Once convergence is achieved, the optimized *A(q)* and *B(q)* are used to calculate a final set of MSDs, which are averaged to obtain the representative MSD of the sample.

### Viscosity calculation from MSD

From the MSD, the diffusion coefficient (D) is obtained from a linear fit of the first four points of MSD versus lag time using:

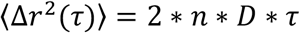

The cytoplasmic viscosity (*η*) is calculated via the Stokes-Einstein relation for spherical particles in a viscous liquid:

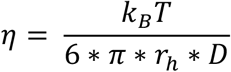

where *k*_*B*_ is the Boltzmann constant, *T* the absolute temperature, and *r*_*h*_ the particle radius.

### Statistical Analysis

At least 30 cells per condition were analyzed. Outliers were detected using MATLAB’s *rmoutlier* function and defined as elements more than 3 standard deviations from the mean. They were removed when corresponding to blurry or unhealthy cells. Statistical significance was assessed using Welch’s ANOVA using GraphPad Prism 10 software.

## Supplementary information

**Figure S1.**
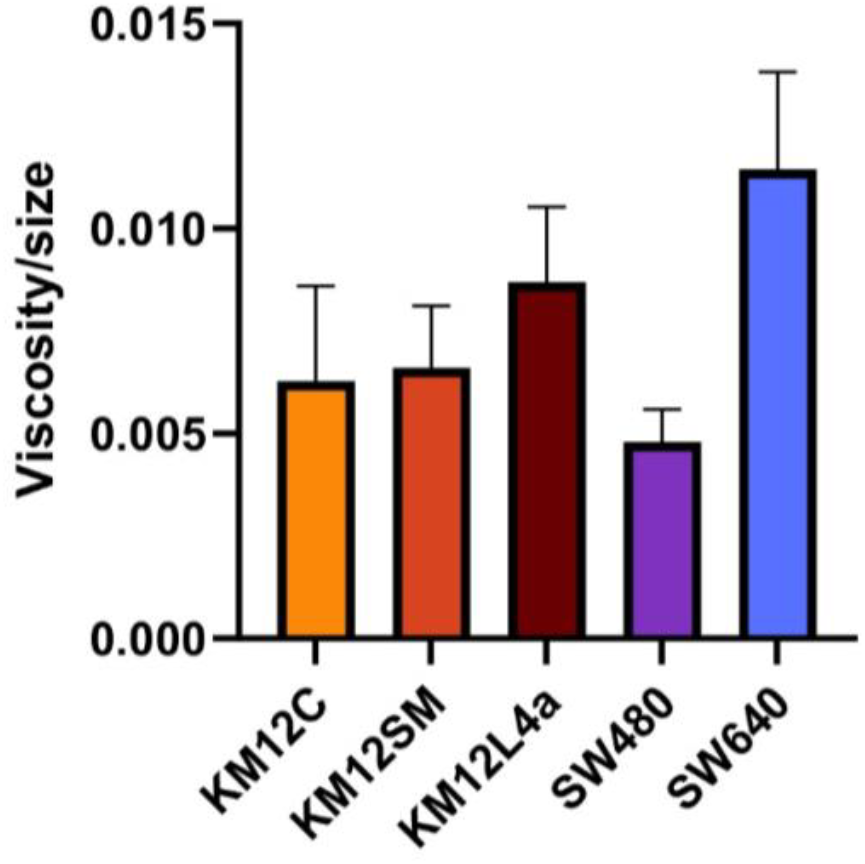
Average viscosity per cell line, normalized by cell area. Plotted as geometric mean with 95% confidence interval, as all datasets showed a lognormal distribution. Number of cells n = 25, 24, 27, 28, 24 for KM12C, KM12SM, KM12L4a, SW480 and SW620 respectively.

## References

Aseervatham, J. (2020). Cytoskeletal Remodeling in Cancer. Biology, 9(11), 385. 10.3390/BIOLOGY9110385

Berret, J.-F. (2015). Local viscoelasticity of living cells measured by rotation magnetic spectroscopymagnetic spectroscopy. Nature Communications, 7(10134). 10.1038/ncomms10134

Camps, J., Morales, C., Prat, E., Ribas, M., Capellá, G., Egozcue, J., Peinado, M. A., & Miró, R. (2004). Genetic evolution in colon cancer KM12 cells and metastatic derivates. International Journal of Cancer, 110(6), 869–874. 10.1002/IJC.20195;PAGE:STRING:ARTICLE/CHAPTER

Cerbino, R., & Cicuta, P. (2017). Perspective: Differential dynamic microscopy extracts multi-scale activity in complex fluids and biological systems. Journal of Chemical Physics, 147(11), 110901. 10.1063/1.5001027/1062993

Cerbino, R., & Trappe, V. (2008). Differential dynamic microscopy: Probing wave vector dependent dynamics with a microscope. Physical Review Letters, 100(18), 188102. 10.1103/PHYSREVLETT.100.188102/CERBINO_420NM.MOV

Datta, A., Deng, S., Gopal, V., Yap, K. C. H., Halim, C. E., Lye, M. L., Ong, M. S., Tan, T. Z., Sethi, G., Hooi, S. C., Kumar, A. P., & Yap, C. T. (2021). Cytoskeletal Dynamics in Epithelial-Mesenchymal Transition: Insights into Therapeutic Targets for Cancer Metastasis. Cancers 2021, Vol. 13, Page 1882, 13(8), 1882. 10.3390/CANCERS13081882

Delarue, M., Brittingham, G. P., Pfeffer, S., Surovtsev, I. V., Pinglay, S., Kennedy, K. J., Schaffer, M., Gutierrez, J. I., Sang, D., Poterewicz, G., Chung, J. K., Plitzko, J. M., Groves, J. T., Jacobs-Wagner, C., Engel, B. D., & Holt, L. J. (2018). mTORC1 Controls Phase Separation and the Biophysical Properties of the Cytoplasm by Tuning Crowding. Cell, 174(2), 338-349.e20. 10.1016/J.CELL.2018.05.042

Dessard, M., Manneville, J. B., & Berret, J. F. (2024). Cytoplasmic viscosity is a potential biomarker for metastatic breast cancer cells. Nanoscale Advances, 6(6), 1727–1738. 10.1039/D4NA00003J

Edera, P., Bergamini, D., Trappe, V., Giavazzi, F., & Cerbino, R. (2017). Differential dynamic microscopy microrheology of soft materials: A tracking-free determination of the frequency-dependent loss and storage moduli. Physical Review Materials, 1(7), 073804. 10.1103/PhysRevMaterials.1.073804

Guigas, G., Kalla, C., & Weiss, M. (2007). Probing the nanoscale viscoelasticity of intracellular fluids in living cells. Biophysical Journal, 93(1), 316–323. 10.1529/biophysj.106.099267

Hernandez, C. M., Duran-Chaparro, D. C., van Eeuwen, T., Rout, M. P., & Holt, L. J. (2024). Development and Characterization of 50 nanometer diameter Genetically Encoded Multimeric Nanoparticles. BioRxiv : The Preprint Server for Biology. 10.1101/2024.07.05.602291

Hewitt, R. E., Mcmarlin, A., Kleiner, D., Wersto, R., Martin, P., Tsoskas, M., Stamp, G. W. H., Stetler-Stevenson, W. G., & Hewitt, R. (n.d.). Validation of a model of colon cancer progression. 10.1002/1096-9896(2000)9999:9999

Keshmiri, H., Cikes, D., Samalova, M., Schindler, L., Appel, L.-M., Urbanek, M., Yudushkin, I., Slade, D., Weninger, W. J., Peaucelle, A., Penninger, J., & Elsayad, K. (2023). Imaging the microscopic viscoelastic anisotropy in living cells. 10.1101/2023.05.28.542585

Korunova, E., Sikirzhystki, V., Twiss, J. L., Vasquez, P., & Shtutman, M. (2024). Single Particle Tracking of Genetically Encoded Nanoparticles: Optimizing Expression for Cytoplasmic Diffusion Studies. BioRxiv, 2024.11.17.623896. 10.1101/2024.11.17.623896

Kuimova, M. K., Botchway, S. W., Parker, A. W., Balaz, M., Collins, H. A., Anderson, H. L., Suhling, K., & Ogilby, P. R. (2009). Imaging intracellular viscosity of a single cell during photoinduced cell death. Nature Chemistry 2009 1:1, 1(1), 69–73. 10.1038/nchem.120

Morse, P. D. (1986). [17] Determining intracellular viscosity from the rotational motion of spin labels. Methods in Enzymology, 127(C), 239–249. 10.1016/0076-6879(86)27020-2

Prevedel, R., Diz-Muñoz, A., Ruocco, G., & Antonacci, G. (2019). Brillouin microscopy: an emerging tool for mechanobiology. Nature Methods, 16(10), 969–977. 10.1038/S41592-019-0543-3;SUBJMETA

Srivastava, A., & Krishnamoorthy, G. (1997). Cell Type and Spatial Location Dependence of Cytoplasmic Viscosity Measured by Time-Resolved Fluorescence Microscopy. Archives of Biochemistry and Biophysics, 340(2), 159–167. 10.1006/ABBI.1997.9910

Verwei, H. N., Lee, G., Leech, G., Petitjean, I. I., Koenderink, G. H., Robertson-Anderson, R. M., & McGorty, R. J. (2022). Quantifying Cytoskeleton Dynamics Using Differential Dynamic Microscopy. Journal of Visualized Experiments : JoVE, 2022(184). 10.3791/63931

Wiseman, P. W. (2013). Image Correlation Spectroscopy: Mapping Correlations in Space, Time, and Reciprocal Space. Methods in Enzymology, 518, 245–267. 10.1016/B978-0-12-388422-0.00010-8

